# Accessible and fast single-cell proteomics by DIA integrating Tecan UNO cell dispensing platform, Vanquish Neo LC and Exploris 480 MS

**DOI:** 10.1101/2025.10.01.679734

**Authors:** Amanda Suárez-Fernández, Judit Bestilleiro-Márquez, Gonzalo Sánchez-Duffhues, Ignacio Ortea

## Abstract

Recent technological advancements in high-sensitivity LC-MS platforms and MS acquisition strategies have addressed key analytical challenges in single-cell proteomics (SCP), paving the way for SCP to become a transformative tool in biology and medicine. Techniques such as MS2 DIA and HRMS1 DIA exploit wide isolation windows and high-resolution MS1 and MS2 spectra to maximize protein identifications, even from low-abundance signals. In parallel, improved data processing algorithms reduce missing values and enhance the reliability of SCP data. These advancements are complemented by automated cell isolation technologies, which ensure precise single-cell handling with minimal sample loss, and in some implementations enable integrated, automated downstream sample preparation steps.

Despite this progress, practical, end-to-end best practices for MS-based SCP remain an active area of optimization. Severe sample limitation, adsorption-related losses, the need for robust quantification across hundreds to thousands of single cells, and the lack of comprehensive protocols that systematically integrate and compare critical workflow variables complicate method selection and SCP implementation.

In this context, this study builds on recent LC-MS based SCP advancements by presenting a complete, optimized, minimum user operation SCP workflow that integrates the Tecan UNO cell dispensing platform, a Vanquish Neo LC system, and an Orbitrap Exploris 480 mass spectrometer, applied to human primary cells. The workflow is designed to minimize data/cell loss, ensure high reproducibility, and maximize identifications. We evaluate key parameters, including acquisition schemes and LC gradient and column, and distill practical recommendations. The resulting protocol provides a foundation for an easy and accessible routine SCP application in biology and medicine.

## 1. Introduction

Single-cell proteomics (SCP) by LC-MS, which has emerged as a transformative approach in the field of proteomics, enables direct quantification of protein expression at the resolution of individual cells, resolving the cellular heterogeneity that underlies diverse biological processes such as development, differentiation, and disease. In contrast, traditional bulk proteomics, while powerful, average the proteomic profiles across large cell populations, diluting the unique characteristics of individual cells and limiting insights into rare cell populations or dynamic cellular states. Cell functional states, differentiation trajectories, and pathological mechanisms previously inaccessible can now be revealed using single cell analysis of protein expression [1– 3]. Its potential is reflected in high-impact biomedical applications such as oncology, immunology, and neuroscience, where cell-to-cell variability shapes disease course and treatment response, improves detection of off-target effects, refines dose-response assessment, and informs biomarker and target discovery to guide patient-personalized therapeutic strategies [4–9].

In recent years, methodological advances have addressed key challenges in every stage of the SCP workflow (Figure 1), pushing the field from proof-of-concept demonstrations to discovery and hypothesis-driven studies. Innovations include miniaturization and automation of cell isolation and sample preparation to reduce yield loss, advances in low-nanoflow LC, gas-phase ion separation (e.g. FAIMS), highly sensitive MS acquisition strategies, and improved data processing and analysis pipelines that increase identification depth, quantitative accuracy, and data completeness [10–12].

**Figure 1.**
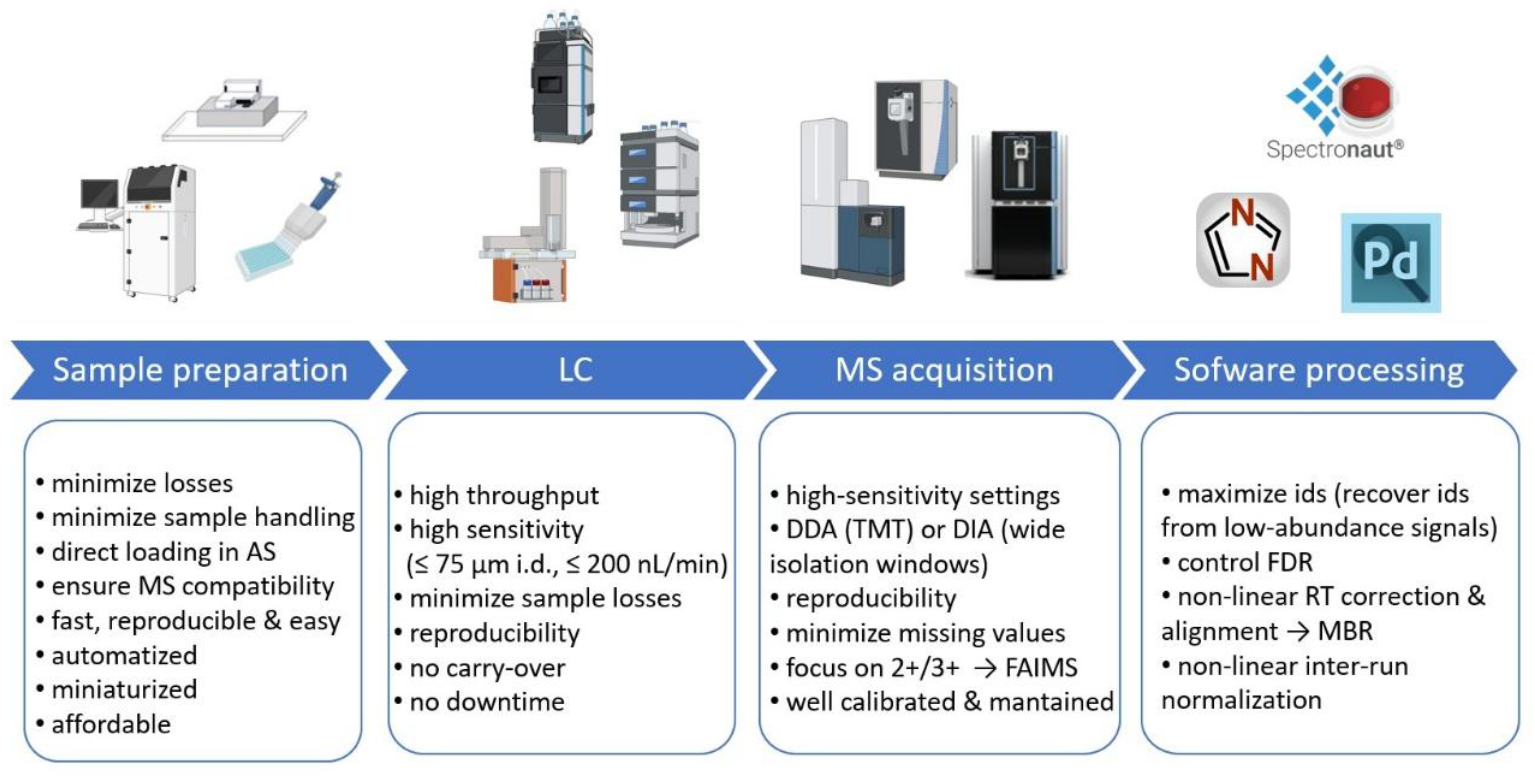
Key challenges in single-cell proteomics. AS, autosampler; i.d., internal diameter; DDA, data-dependent acquisition; TMT, tandem mass tag; DIA, data-independent acquisition; FAIMS, field asymmetric ion mobility spectrometry; ids, identifications; FDR, false discovery rate; RT, retention time; MBR, match-between-runs.

High-fidelity dispensing of single cells and sample-preparation reagents is foundational to any SCP workflow. Although conventional cell sorters are used to isolate single cells, they often exhibit bias towards different cell types and typically require subsequent off-line steps for cell lysis and protein digestion, increasing handling and potential losses. The cellenONE platform, a commercially available system that integrates single-cell dispensing with on-chip, one-pot sample preparation, addresses this limitation by avoiding manual transfers, and has shown improved performance, becoming a prevalent solution for SCP sample preparation [13,14]. Notably, Sanchez-Avila *et al*. demonstrated that the Tecan UNO, a cost-efficient, thermal-inkjet based cell and reagent dispenser, enables rapid, one-step sample preparation. This platform allows high single-cell confirmation rates and accurate delivery of volumes down to pL to low-μL into 96- and 384-well standard plates [15]. Reported results are promising, with 2572 and 1668 proteins identified from individual HeLa cells using a 30-μm I.D. home-packed column and a commercial PepMap 100 (Thermo) column, respectively.

On the chromatographic separation side, sensitivity and throughput are governed by column geometry and flow regime. High-sensitivity, low-nanoflow LC configurations using narrow-bore columns (e.g. 30-50 μm I.D.) operated at 50-150 nL min^−1^ have proven particularly effective for sample-limited proteomics, delivering deeper proteome coverage at practical samples per day (SPD) rates. Tuning of column I.D., gradient length, and loading/injection strategy balances identification depth with robustness for large SCP series, providing an empirical basis for method selection in SCP workflows [16].

Equally important are acquisition strategies that maximize identifications and data completeness at picogram inputs. Early SCP workflows often relied on isobaric labeling with tandem mass tag (TMT) using data-dependent acquisition (DDA), typically with a carrier channel, to multiplex tens of single cells per run and minimize analysis time per cell. With higher sensitivity instruments and optimized low-flow LC, label-free data-independent acquisition (DIA) can now achieve deeper per-cell proteome coverage, data completeness, and between-run consistency [11]. Therefore, sensitivity-tailored DIA has emerged as a strong option for low-input and single-cell analyses. DIA further synergizes with modern computational frameworks to limit interferences and strengthen peptide- and protein-level quantification across large cohorts [17]. Although label-free DIA quantification is conventionally performed at the fragment-ion (MS2) level, DIA also acquires MS1 spectra, and leading software packages (e.g., Spectronaut, DIA-NN) support quantification at both levels. In the HRMS1-DIA scheme, the precursor m/z range covered by the MS2 isolation windows is partitioned into segments, with higher resolution MS1 scans interleaved between these segments [17]. Using these MS1 scans for quantification increases points-per-peak (PPP) and allows higher Orbitrap resolution and longer injection times. Petrosius et al. [17] combined wide MS2 isolation windows with HRMS1-based quantification in a method termed WISH-DIA (Wide Isolation-window High-resolution MS1-DIA).

Advances in instrumentation continue to expand achievable depth and quantitative precision. For example, the Orbitrap Astral mass spectrometer combines very high scan speed with improved ion transmission, enabling deeper coverage for low-input and single-cell samples. Method evaluations on Astral platforms indicate that DIA parameterization (e.g., isolation window schemes, injection times, cycle time, and scan scheduling) materially impacts sensitivity in the single-cell range, underscoring the need for systematic benchmarking when translating SCP methods across instruments [18].

Despite such progress, practical best practices for end-to-end MS-based SCP remain under active optimization. Ultralow sample amounts, susceptibility to adsorption- and transfer-related losses, and the need for robust quantification across hundreds to thousands of single cells complicate method design. Moreover, comprehensive protocols that systematically integrate and compare key workflow variables remain limited [11,17]. Optimizing individual components, such as column geometry or gradient design, does not necessarily translate into gains at the level of the entire workflow.

Our study addresses this gap through the optimization and thorough description of a complete and reproducible label-free SCP protocol covering: (i) single-cell dispensing; (ii) low-loss, one-pot cell lysis and protein digestion; (iii) LC separation using commercial available columns and short-to-medium gradients on standard setups; (iv) comparative evaluation of DIA variants on a Q-Orbitrap platform; and (v) bioinformatic processing with and without external high-concentration reference runs to assess the impact on identifications and quantitative stability. Across these dimensions, we benchmark performance and contextualize gains relative to recent SCP literature, including those employing the latest high-end mass spectrometers, which offer nominally higher performance at substantially greater cost. By integrating and contrasting multiple critical parameters within a single framework, this work not only complements the most influential recent contributions, but provides a practical, reproducible guide for laboratories seeking robust, scalable, and accessible label-free SCP measurements that could be generalized across cell types and experimental designs. Because the protocol minimizes user intervention and relies on commercially available hardware and standard, unmodified setups (without in-house or custom modifications), we are confident that it lowers barriers to adoption for general proteomics groups and researchers, including first-time users, and even enables straightforward deployment in core facilities.

## 2. Materials and methods

### 2.1. Cells

Bone marrow-derived human mesenchymal stem cells (hMSCs) have been described before [19]. The cells were cultured in Alpha MEM (Gibco, cat. 32561094) supplemented with 1 ng/ml bFGF (Sigma, cat. F0291), 10% FBS and Pen/Strep. The cells were cultured in a CO_2_ controlled 37 °C incubator. Cell passaging was performed using standard EDTA/Trypsin incubation. For sample preparation, the cells were trypsinized, counted and resuspended in phosphate buffered solution (PBS) at a concentration of 10^5^ cells/mL.

### 2.2. Single-cell dispensing and sample preparation

Single-cell samples were prepared on the UNO platform (Tecan) using the instrument control software (UnoControl; HP Development Company, L.P.), following a previously published protocol with minor modifications [15]. In a first step, individual cells were dispensed into 96-well plates using cell-specific C1a cassettes (Tecan). Given a general size of 15-30 μm for human MSCs, the size gate was set to Extra Large (21–25 μm), with the fluid group set to “Cells.” In a second step, one-pot cell lysis and protein digestion were performed by dispensing 500 nL per well of a lysis/digestion cocktail (1 ng trypsin and 1 ng Lys-C in 0.05% n-dodecyl-β-D-maltoside, DDM) using D1 cassettes (Tecan) and the “Aqueous + Brij 35” fluid class setting. Negative control wells (with lysis/digestion cocktail but no cell) and blank wells (no cell and no cocktail) were included. Plates were covered with a silicone sealing mat and incubated at 70 °C for 1 h, cooled down to 4 °C for 15 min, centrifuged at 600 × g for 1 min, sealed with aluminum film, and stored at –20 °C until analysis.

### 2.3. LC-MS analysis

Immediately prior to LC–MS analysis, the aluminum sealing foil was removed and 4 μL of solvent A (0.1% formic acid in water) was dispensed into each well using the Tecan UNO. Plates were then covered with a pre-pierced plastic cap mat (Thermo Fisher Scientific), following the manufacturer’s recommendations, and placed directly in the autosampler of the Vanquish Neo LC system (Thermo Fisher Scientific). To ensure complete transfer of the well contents, the injection volume was set to 4.5 μL per well.

Two commercial columns were evaluated in a standard trap-and-elute configuration: PepMap Neo, 75 μm × 15 cm (Thermo Fisher Scientific), and Aurora Elite, 75 μm × 15 cm (IonOpticks). Three LC gradients were tested (Figure 2A): a 25 min gradient (36 min total runtime; 40 SPD) for PepMap Neo, and 24 min and 14.5 min gradients (35 min and 21 min total runtime; 40 and 70 SPD, respectively) for Aurora Elite. Importantly, solvent B composition differed by column, according to manufacturer guidance: 80% acetonitrile (ACN) with 0.1% formic acid (FA) for PepMap Neo, and 100% ACN with 0.1% FA for Aurora Elite. Solvent A was water with 0.1% FA for both columns. Gradients were run at 200 nL/min; to accelerate peptide elution on the Aurora Elite setup, the first 2 min were run at 400 nL/min, and the final column-wash step in the 70 SPD method was also executed at 400 nL/min to shorten the total runtime (Figure 2A).

**Figure 2.**
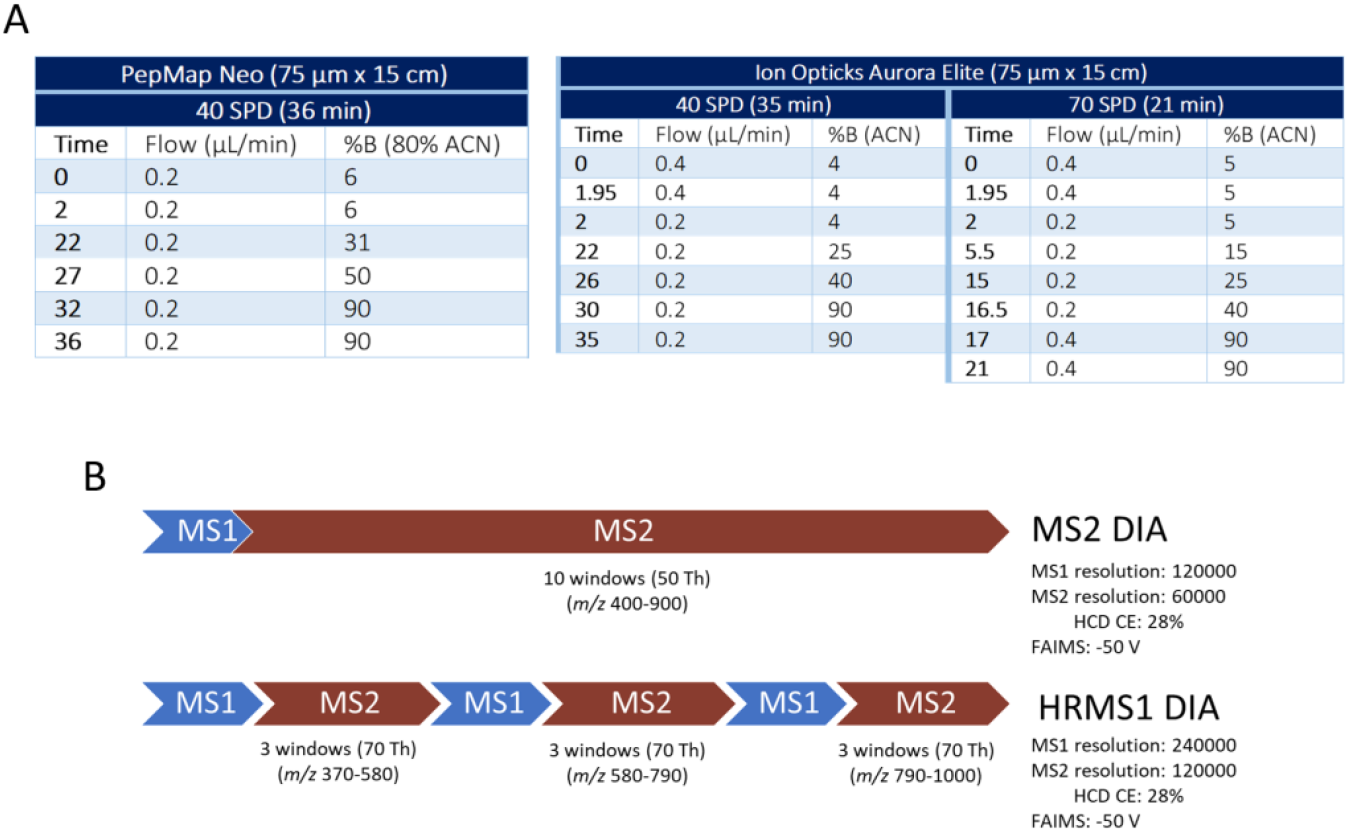
Optimized LC-MS methods used. (A) LC column and gradient combinations. (B) DIA schemes.

Analyses were performed on an Orbitrap Exploris 480 mass spectrometer (Thermo Fisher Scientific) equipped with a FAIMS Pro interface in DIA mode. Two DIA schemes were optimized (Figure 2B). In the first, MS2-DIA (MS2-based quantification), each cycle comprised one MS1 survey scan (120,000 resolution) followed by 10 MS2 windows of 50 Th spanning the m/z range 400-900 (Orbitrap resolution 60,000). In the second, HRMS1-DIA (MS1-based quantification), the precursor range m/z 370-1000 was divided into three segments (m/z 370-580, 580-790, and 790-1000); within each segment, three wider MS2 windows (70 Th) were acquired, with high-resolution MS1 scans interleaved between segments (Figure 2B). For HRMS1-DIA, MS1 and MS2 resolutions were 240,000 and 120,000, respectively. In both methods, normalized HCD collision energy was 28%, and compensation voltage was set to -50 V at the FAIMS Pro interface for ion mobility filtering.

### 2.4. Data Processing

Raw DIA data were analyzed in Spectronaut v19.0 (Biognosys) using the directDIA workflow. Searches (Spectronaut’s Pulsar engine) used a human canonical proteome database (UniProtKB/Swiss-Prot; downloaded May 10, 2024) concatenated with the MaxQuant common contaminants list. Spectronaut factory settings were applied, with the following exceptions: proteases, Trypsin/P and Lys-C; variable modifications, protein N-terminal acetylation and methionine oxidation; fixed modifications, none. False discovery rates were controlled to <1% at the PSM, peptide, and protein-group levels.

Cross-run normalization was kept at Automatic (Local for datasets <500 runs). Missing values were not imputed (default in Spectronaut). Protein label-free quantification method was kept at Automatic (MaxLFQ algorithm for datasets <500 runs). The Quantity MS Level parameter remained at MS2 (all qualifying fragment ions) for MS2-DIA, and was changed to MS1 (monoisotopic precursor plus isotopic envelope) for HRMS1-DIA. We also evaluated the hybrid library extension option in Spectronaut by adding three high-concentration reference runs (10 ng K562 commercial digest) as library-extension runs.

### 2.5. Pathway and Gene Ontology (GO) analysis

To assess the coverage of the human proteome captured by the optimized workflow, quantified protein groups were mapped to Reactome biological pathways (https://reactome.org/, accessed 27 August 2025) [20]. To examine coverage within specific pathways, the quantified protein groups were analyzed with ShinyGO v0.82 (https://bioinformatics.sdstate.edu/go82/, accessed 27 August 2025) [21] using the KEGG pathway maps database [22].

## 3. Results and discussion

First, we processed hMSCs on the Tecan UNO to optimize single cell dispensing and on-plate sample preparation, then analyzed these true single cell samples by LC-MS DIA to assess performance and to detect and correct practical issues. Importantly, consumables proved critical, particularly the microplates and sealing film/mats. Plate choice is important for reducing the binding and loss of proteins and peptides during sample processing, and it is critical that they are compatible with the LC autosampler, in order to prevent liquid transfer to vials. We used Eppendorf 96-well Lo-bind plates with consistent results, whereas alternative plates produced markedly fewer quantified proteins (*data not shown*). Sealing materials also mattered. Depending on composition, some seals can shed particulates that obstruct the Vanquish Neo sample seat. This occurred with the aluminum sealing film described in the protocol by Sanchez-Avila et al. [15], but not with pre-pierced plastic cap mats (Thermo Fisher Scientific).

These initial real single cell samples were also used to optimize the combination of LC gradients and DIA method. We evaluated three gradients using two commercial 75 um i.d. analytical columns designed for high-sensitivity and standard nanoLC-MS flow rates. To enable a direct comparison of both columns, we adjusted the 40 SPD gradient to the organic composition of solvent B used for each particular column (80% ACN for PepMap Neo, 100% for Aurora Elite), so that both 40 SPD methods were equivalent in ACN profile. For higher throughput (70 SPD), we did not simply shorten the gradient; instead, we re-shaped the slope, reducing steepness in the peptide-dense region to preserve identifications while decreasing runtime. All experiments used a trap-and-elute setup. Transfer-induced losses during sample preparation are well documented [17], and therefore it is important to use chromatography systems that allow direct loading from the plate on which the single cell samples are processed, eliminating intermediate vial transfers. Because the samples are loaded without off-line cleanup after lysis and digestion, we favored a trap-and-elute configuration over direct injection. Additionally, trap columns are often preferred over direct injection in SCP because they accelerate sample loading by allowing high-flow transfer of samples to the analytical column [11].

### 3.1. Performance on low-abundance samples

To evaluate performance, we analyzed low-abundance K562 digests at 10 ng and 400 pg using three column–gradient combinations and two acquisition schemes (MS2-DIA and HRMS1-DIA). We also assessed the impact of using a hybrid library extension by adding three 10-ng K562 runs as extension runs. The largest variation in quantified protein groups was attributable to column choice. At 40 SPD, moving from PepMap Neo to Aurora Elite increased identifications from approx. 4000 to 6000 at 10 ng and from a approx. 3000 to 4000 at 400 pg (Figure 3A). Increasing throughput on Aurora to 70 SPD reduced identifications, but still slightly exceeded those obtained with PepMap Neo with a longer gradient. Regarding MS acquisition, for the 10 ng samples, MS2-DIA slightly outperformed HRMS1-DIA in terms of protein identifications across LC conditions. However, that was not the case at 400 pg, since both schemes yielded similar counts except for Aurora 40 SPD, where HRMS1-DIA showed higher sensitivity. Enabling the hybrid library extension clearly increased identifications for the 400-pg runs across all LC and acquisition combinations assessed.

**Figure 3.**
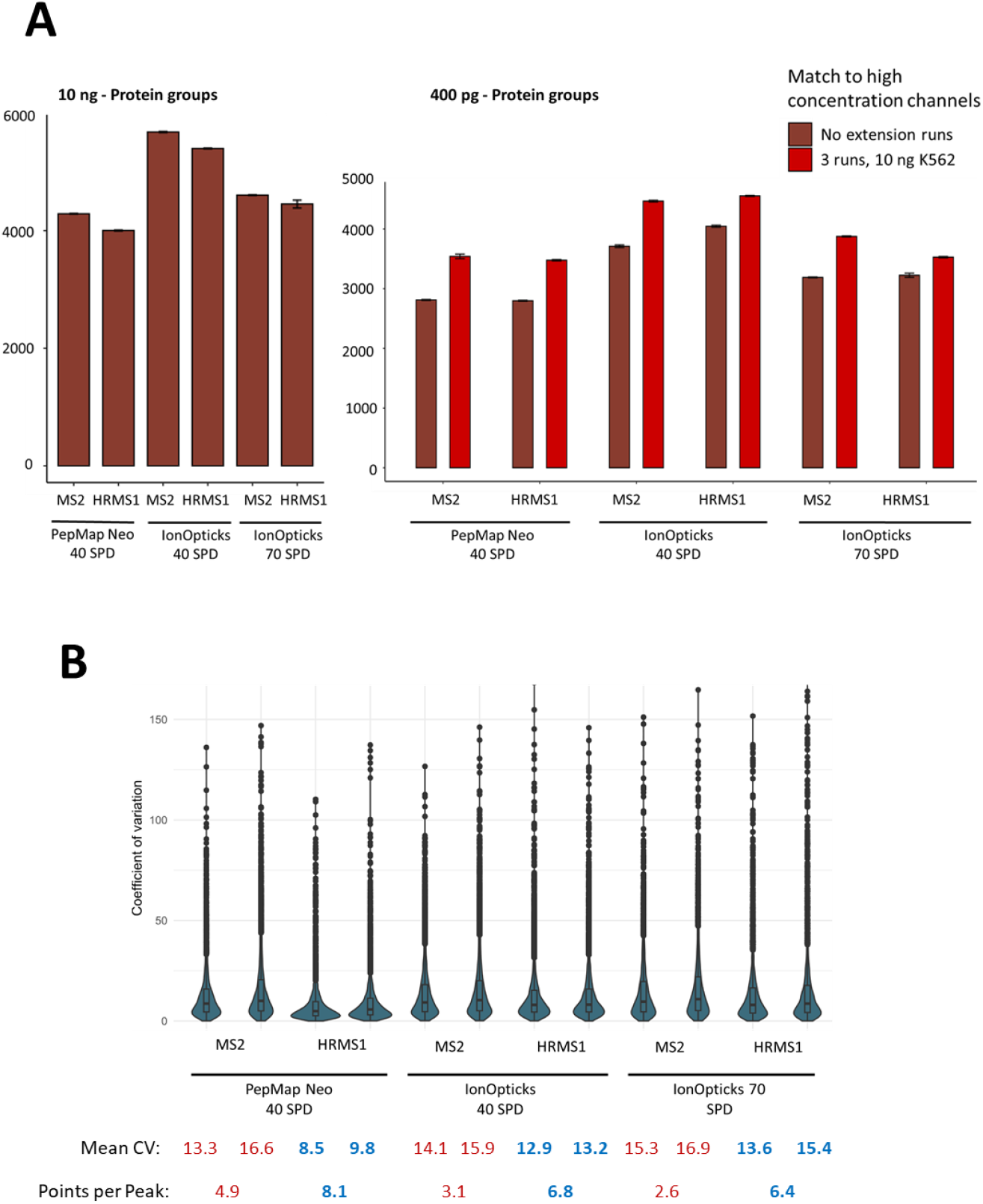
Performance of K562 digests at 10 ng and 400 pg across the tested workflows. We evaluated three column–gradient combinations, two DIA acquisition schemes (MS2-DIA and HRMS1-DIA), and the effect of using a hybrid library extension. (A) Protein groups quantified per workflow (mean ± SD; n = 3). (B) Coefficient of variation (CV, %) and mean points-per-peak (PPP) per workflow for the 400 pg K562 injections (n = 3). For each instrumental workflow, bars are shown in pairs: left, without extension runs; right, with Hybrid Library Extension.

Reproducibility was maintained across conditions, with mean coefficients of variation (CVs) below 17% (Figure 3B), although notable differences emerged by workflow. PepMap Neo delivered the lowest CVs despite fewer protein groups, consistent with its broader chromatographic peaks that yield higher points-per-peak (PPP) and thus improved quantitative precision at the expense of sensitivity. HRMS1-DIA, by increasing PPP, further reduced mean CVs, most prominently with PepMap Neo. Switching from 40 to 70 SPD on Aurora had a minor effect on precision, with a slight CV increase accompanying a small decrease in PPP. In contrast, enabling the hybrid library extension modestly worsened CVs. We hypothesize this may reflect increased introduction of false positive matches during identification. Overall, for the 400 pg injections, the highest reproducibility was achieved with PepMap Neo at 40 SPD using HRMS1-DIA and without extension runs.

### 3.2. Protein quantification in real single-cell samples

To assess performance on real samples, hMSC single cells prepared on the Tecan UNO were analyzed under three LC-gradient conditions and two DIA schemes (MS2-DIA and HRMS1-DIA). We also evaluated the use of hybrid library extension enabling the “extension runs” option in Spectronaut.

As Figure 4 shows, at 40 SPD, the Aurora Elite column outperformed PepMap Neo, quantifying >3,000 protein groups (up to 3,528 in total). Increasing throughput to 70-SPD on Aurora reduced identifications relative to Aurora 40 SPD, yet counts remained slightly above those obtained with PepMap Neo at 40 SPD. We also compare HRMS1-DIA and MS2-DIA. In previous studies HRMS1-DIA outperformed standard MS2-DIA, yielding higher identifications and improved quantitative precision [17,23]. However, in our data, HRMS1-DIA performed better with the PepMap Neo configuration but worse with Aurora Elite, underscoring the need to co-optimize LC conditions and MS acquisition parameters for SCP rather than adopt a one-size-fits-all approach.

**Figure 4.**
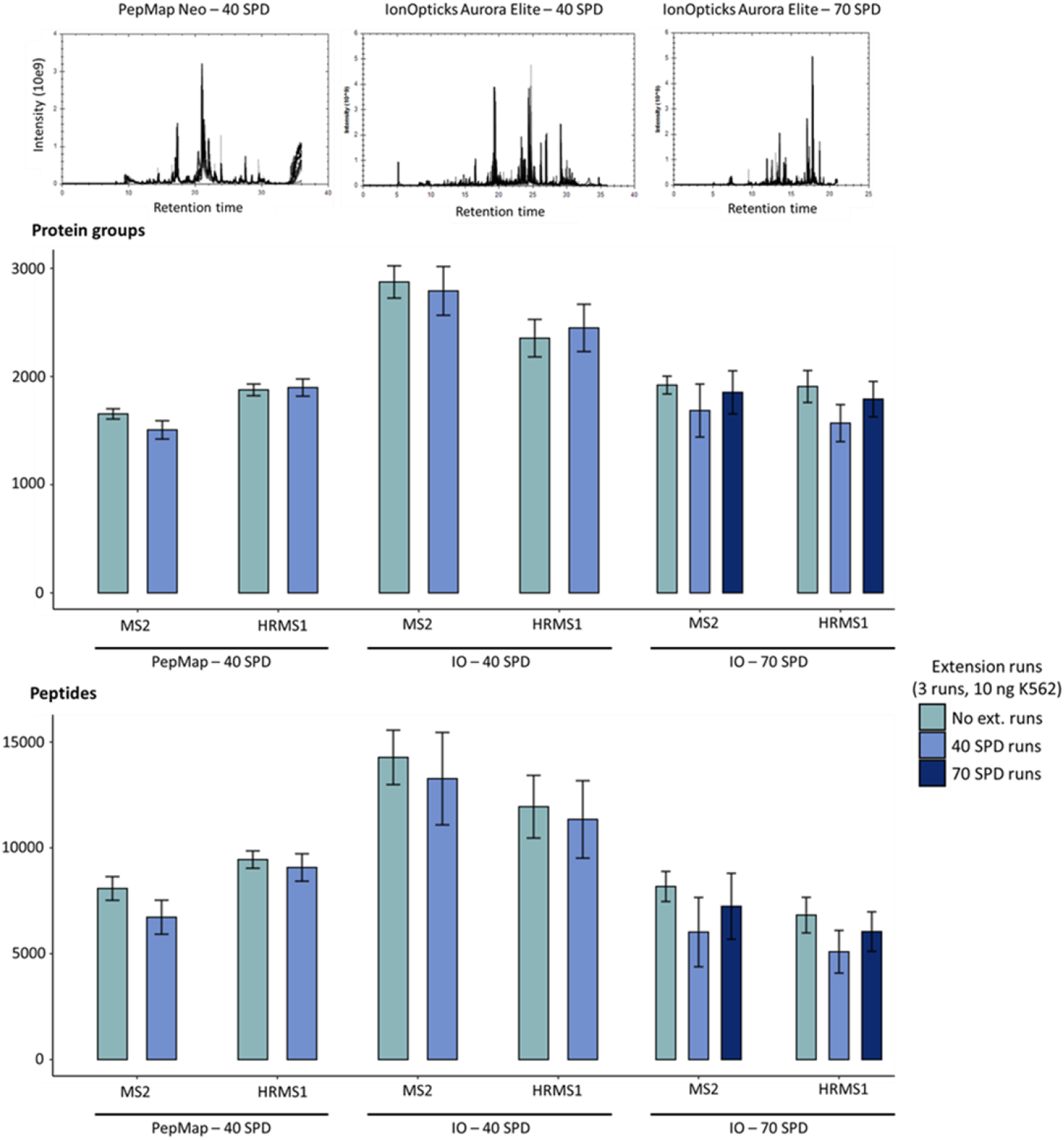
Performance of the tested workflows for hMSCs processed on the Tecan UNO. Three column-gradient combinations, two DIA acquisition schemes (MS2-DIA and HRMS1-DIA), and the effect of enabling Spectronaut Hybrid Library Extension were evaluated. Bars show protein groups and peptides quantified (mean ± SD; n = 25), alongside representative total-ion chromatograms (TICs).

A current strategy to improve identification numbers in spectral library-free approaches is to search alongside matched samples with higher quantities, which are subsequently used to build a hybrid library. In some cases, this approach has shown to improve the number of peptide and protein identifications by >20%, although its precise effects, depending on the number of single cell files included in the search and the specific LC conditions and acquisition scheme, remain to be fully understood [24]. Systematic benchmarking is therefore recommended. When we assessed the use of hybrid library extension, in contrast to the benefit observed for 400 pg K562 digests, real single-cell data showed no improvement, or even a slight decline, in identifications, particularly at the peptide level (Figure 4). Notably, when using the 70 SPD analytical gradient, extension runs acquired with a longer 40 SPD gradient yielded poorer peptide/protein quantification than extension runs acquired with the same 70 SPD gradient. Although longer gradients should, in principle, expand the proteome’s coverage, these results suggest that mismatched gradients likely degrade retention-time alignment and feature transfer, and therefore extension runs should match the analytical gradient to avoid potential issues. Overall, the best performance for hMSCs was obtained without enabling the hybrid library extension.

To contextualize our results, we compared them with recent SCP publications. Ryan Kelly’s group demonstrated that a one-step protease + DDM method identified numbers of proteins comparable to a standard multistep preparation, reaching 1,947 quantified proteins from single HeLa cells prepared on the cellenONE (Cellenion) [25]. When the preparation was ported to the HP D100 (now Tecan UNO), 1,668 proteins were identified from individual HeLa cells using a commercial 50-μm i.d. PepMap 100 column at 100 nL min^−1^, increasing to 2,572 with a 30 μm i.d. home-packed column at 30 nL min^−1^ [15]. Using our optimized workflow and likewise dispensing on the Tecan UNO we achieved higher protein counts on the same mass spectrometer (Orbitrap Exploris 480) while operating under standard nanoLC conditions (commercial 75 μm i.d. columns at 200 nL min^−1^). Zheng et al. [16], using a low-nanoflow LC configuration and an Orbitrap Exploris 480 (with FAIMS interface), reported 3,000 protein groups from 250 pg HeLa digest at 72 SPD and 1,700 from real single HeLa cells at 100 SPD. They used the cellenONE platform for cell isolation and preparation, and a commercial 50 μm i.d. PepMap 100 at 100 nL min^−1^, although connected to the LC (Vanquish Neo) via a low-dispersion Y-piece not broadly available commercially. Despite their lower flow, requiring highly specialized columns, our workflow exceeded those identification numbers while relying on more accessible sample-prep and LC components. Petrosius et al. [17], using 1 ng inputs and the 20 SPD Evosep Whisper method with standard C18 columns, identified ∼1,400 (MS2-DIA) and 2,300 (HRMS1-DIA) protein groups. With a μPAC Neo low-load column (Thermo Fisher Scientific) and a 45-min run (30 SPD), identifications rose to ∼3,000 at 1 ng, decreasing to ∼2,000 at 250 pg. For real single HeLa cells, ∼1,700 (directDIA) and ∼2,000 (with a gas-phase fractionation library) protein groups were reported. Interestingly, and avoiding the need for building a library, we reached above 3,000 protein groups in single hMSCs. On the Astral mass analyzer (Thermo Fisher Scientific), Petrosius et al. [18] identified ∼4,000 and ∼3,000 protein groups with 1 ng and 500 pg injections, respectively. WISH-DIA yielded ∼4,000 at 250 pg, whereas real single-cell datasets ranged ∼1,300 to 3,500 depending on cell size and proteome specialization. Notably, we exceeded those identifications at 400 pg of K562 and quantified up to ∼3,500 protein groups in real single cells, while relying on a more accessible, earlier-generation mass spectrometer with lower performance on paper. With the high-end advances in mass spectrometry instrumentation, incredible proteome coverage has been reached, approaching numbers previously obtainable from bulk proteomics samples. In real single HeLa cells, Ye et al. reported 4,380 (DIA-NN) and 5,204 (Spectronaut) protein groups using the 40 SPD Whisper method on Evosep One, and up to 6,500 with a 60 SPD method on Vanquish Neo LC [24]. Bubis et al. [23] described a coverage of 4,879 proteins in A549 and 3,166 in H460 cells, both lung cancer lines with size ranges similar to hMSCs, ∼20–30 μm. With this instrument configuration (Vanquish Neo LC and Astral mass spectrometer), 5,438 protein groups were reported in 250 pg injections of K562 with a library-free approach, while the use of a tailored library from 10ng of the same digest increased the number of identified protein groups up to 6,800. These studies, approaching ∼5,000 proteins per real single cell, represent a notable SCP depth landmark if widely reproducible. However, in contrast to those setups, our results were obtained with the Tecan UNO dispenser and an Orbitrap Exploris 480, significantly more accessible in a standard lab setting.

When comparing SCP results, a central limitation must be acknowledged: achievable proteome coverage depends strongly on cell size and type. The diversity of cell types and size distributions used across studies makes direct, head-to-head performance assessments challenging. Given differences in cell size and proteome content and complexity, substantial variation in coverage is expected. Beyond size, proteome composition and specialization also influence depth: samples generated from primary material are expected to yield fewer identifications, since more primitive cells (such as multipotency cells) are also expected to have more specialized proteomes [18].

The methods described here remain amenable to further optimization. For example, we did not systematically evaluate the effects of DIA isolation-window width, Orbitrap resolving-power settings, or injection times (ITs). Isolation-window width, in particular, can influence peptide/protein identifications. In the WISH-DIA method optimization by Petrosius et al. [17], the highest identification yields were obtained with 80 m/z windows for MS2-DIA and 40 m/z windows for HRMS1-DIA.

### 3.3. Functional coverage

To evaluate whether the data capture biologically relevant information, we performed pathway and GO analysis using the quantified proteins from real single cell datasets. The optimized SCP workflow resulted in a broad coverage of the human proteome, spanning all major biological pathway groups (Figure 5). Within individual pathways, a substantial fraction of constituent proteins was quantified, as shown in representative examples (Supplementary Figure S1). These examples include key cellular processes such as oxidative phosphorylation, metabolism, protein processing in the endoplasmic reticulum, signal transduction, nucleocytoplasmic transport, and the ribosome. This comprehensive coverage supports the ability of the workflow to deliver biologically meaningful insight into the functional state and key processes of individual cells. Notably, the workflow also showed high sensitivity in detecting proteins involved in disease-associated pathways. Cancer, Alzheimer’s disease, and diabetic cardiomyopathy are shown as examples (Supplementary Figure S2). The broad coverage obtained highlights the potential for applications in disease research and biomarker discovery. The results also emphasize the importance of optimizing LC and MS parameters to maximize pathway coverage and ensure the detection of low-abundance proteins critical for understanding cellular function.

**Figure 5.**
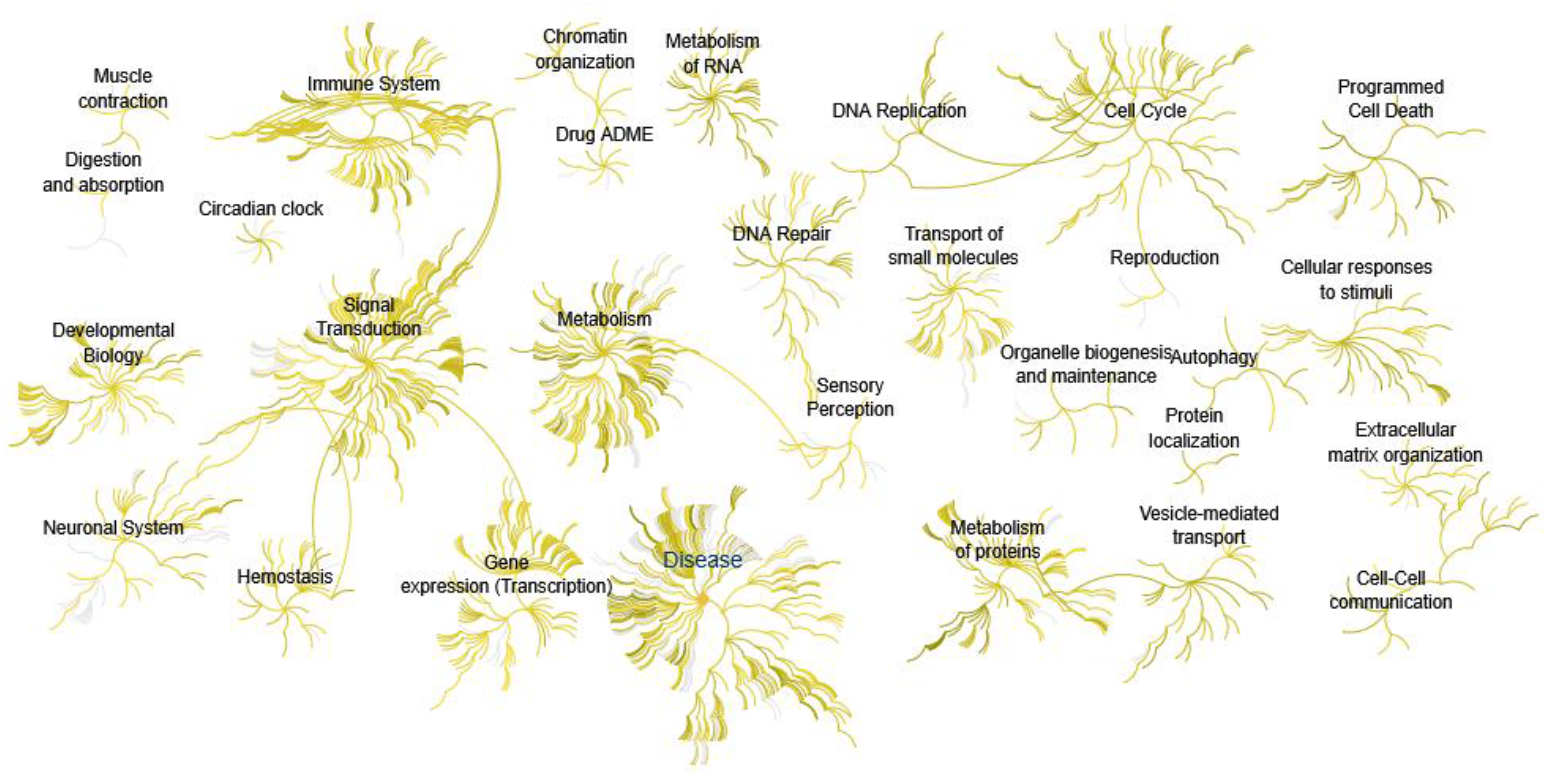
Human-proteome coverage for proteins quantified in hMSCs using the optimized workflow. Each line represents a pathway, grouped by top-level Reactome categories. Pathways containing quantified proteins are highlighted in yellow.

The optimized SCP workflow also provided a broad coverage of GO terms associated with cellular location. Proteins quantified from real single cell datasets mapped to diverse subcellular compartments, including the nucleus, cytoplasm, mitochondria, endoplasmic reticulum, and plasma membrane (Supplementary Figure S3). Notably, coverage also extended to low-abundance organelles such as the Golgi apparatus and lysosomes. These results highlight the ability of the workflow to capture proteins across all cellular locations, providing a holistic view of the proteome within individual cells and supporting biologically relevant insights into cellular organization and function.

## 4. Conclusions

We present a complete, optimized workflow for SCP that integrates the Tecan UNO cell-dispensing platform, a Vanquish Neo LC system, and an Orbitrap Exploris 480 mass spectrometer. The workflow delivers high sensitivity and reproducibility, enabling quantification of up to ∼3,500 protein groups from single human cells. Systematic evaluation of LC gradients, column types, and DIA acquisition schemes highlights the importance of tailoring method parameters to specific experimental needs rather than rely on a single, fixed configuration. The data show broad coverage of cellular components and human pathways, supporting biologically meaningful inferences about cellular organization and function. Notably, the workflow detects low-abundance proteins and achieves functional coverage across disease-relevant pathways, highlighting its utility for applications in disease mechanisms, biomarker discovery, and cellular heterogeneity. Future work will extend the workflow to dissociated tissues and urine-derived single cells, further broadening its applicability and impact in proteomics.

Beyond performance, the workflow emphasizes practical accessibility. The Tecan UNO lowers the barrier to entry for laboratories without bespoke microfluidics and minimizes user intervention through automated dispensing and one-pot preparation. Its compatibility with autosamplers, together with the use of standardized plates and simplified chemistries, supports direct-from-plate injections and reduces manual handling, factors that can accelerate adoption. Consequently, the approach is more accessible to general proteomics groups and amenable to straightforward deployment in core facilities.

## Supplementary Material

**Figure S1**. Coverage within selected pathways representing key biological processes. Quantified proteins are highlighted in red, illustrating the depth achieved for specific routes.

**Figure S2**. Coverage within selected disease-associated pathways. Quantified proteins are highlighted in red, illustrating the depth achieved for specific routes.

**Figure S3**. Gene Ontology-cellular component analysis. Subcellular components to which the quantified proteins were mapped are shown.

