## Supplementary Figures S1-S3 for "Accessible and fast single-cell proteomics by DIA integrating Tecan UNO cell dispensing platform, Vanquish Neo LC and Exploris 480 MS"

### Supplementary Material

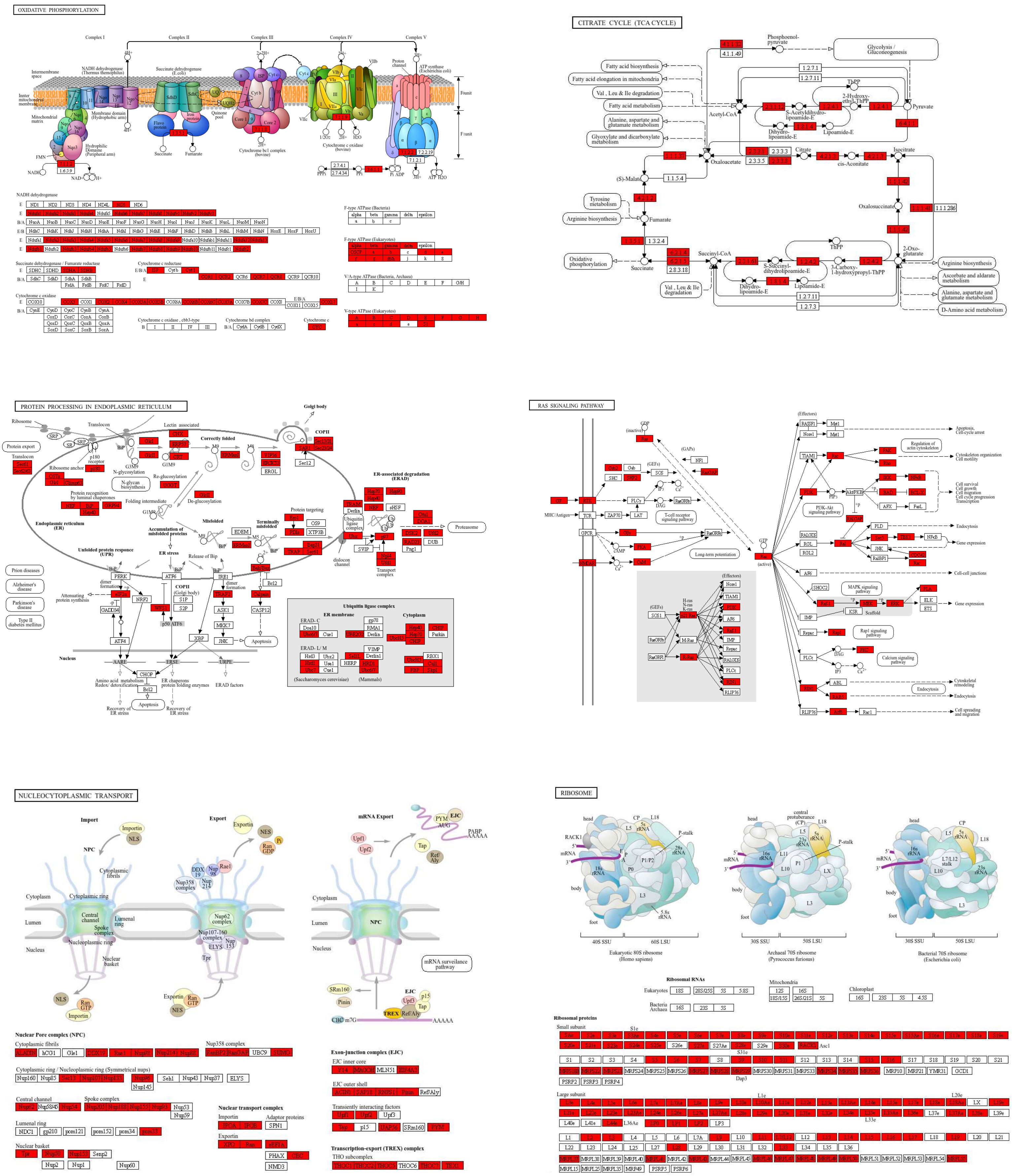


**Figure S1.** Coverage within selected pathways representing key biological processes. Quantified proteins are highlighted in red, illustrating the depth achieved for specific routes.

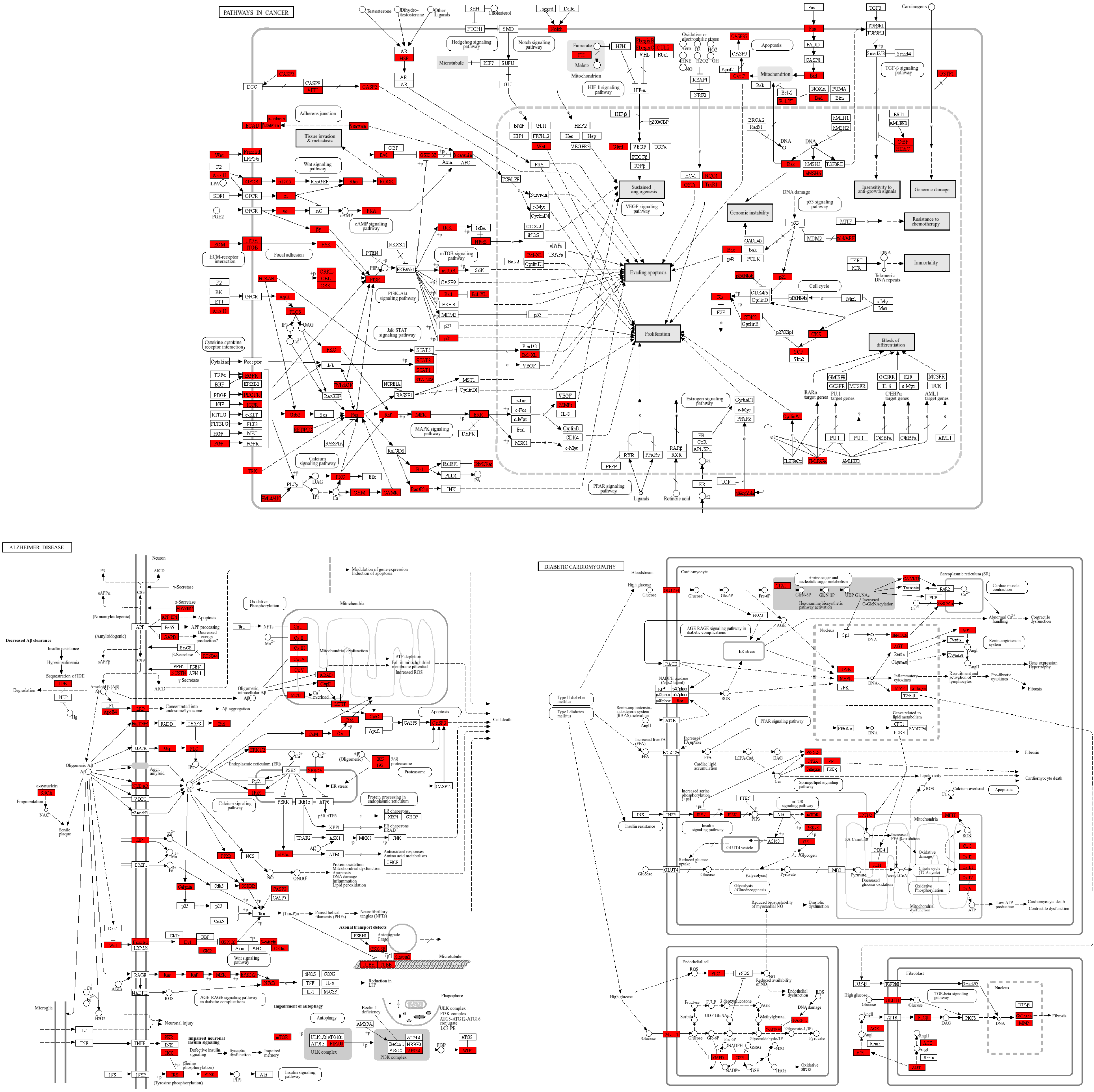


**Figure S2.** Coverage within selected disease-associated pathways. Quantified proteins are highlighted in red , illustrating the depth achieved for specific routes.

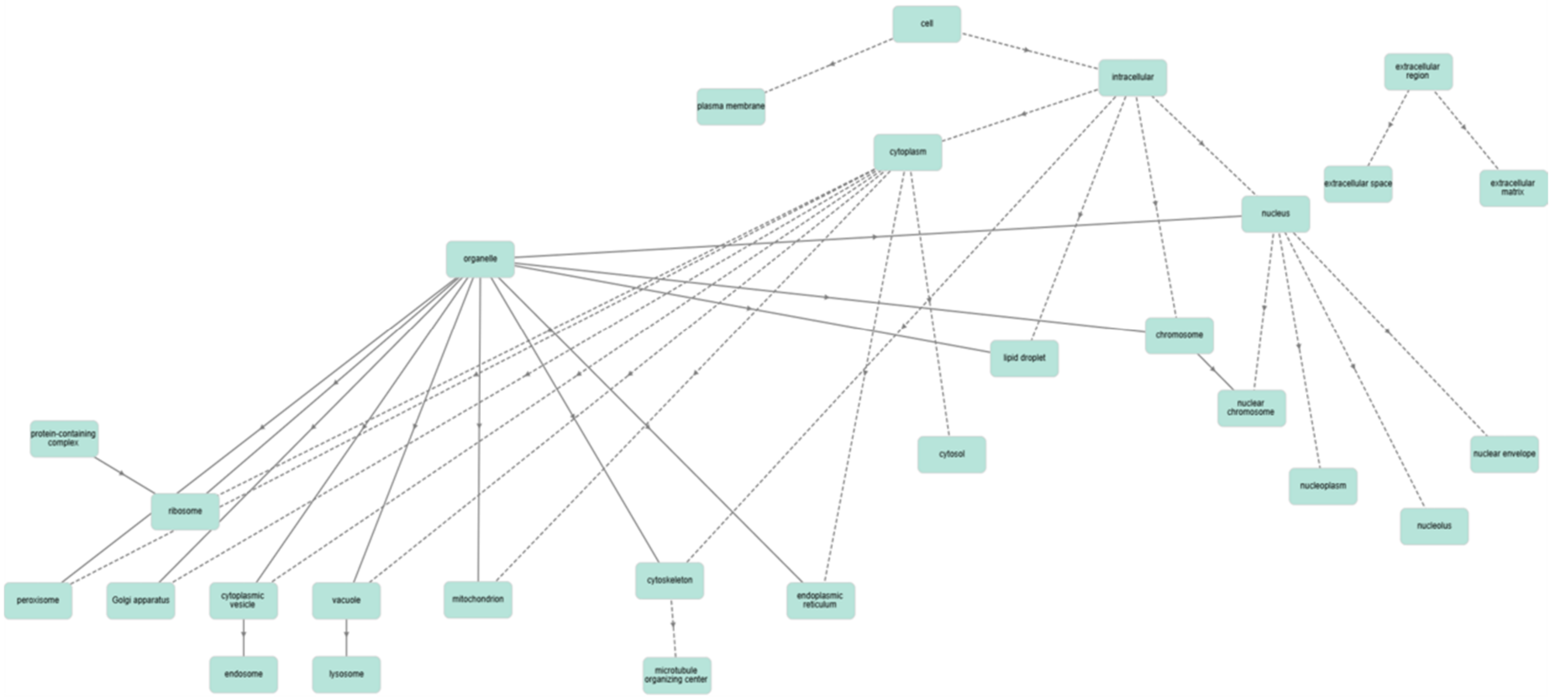


**Figure S3.** Gene Ontology-cellular component analysis. Subcellular components to which the quantified proteins were mapped are shown.
